# *Sinoseris* (Crepidinae, Cichorieae, Asteraceae), a new genus endemic to China of three species, two of them new to science

**DOI:** 10.1101/680843

**Authors:** Ze-Huan Wang, Norbert Kilian, Ya-Ping Chen, Hua Peng

**Author notes:** Authors for correspondence. Norbert Kilian,; Hua Peng.

## Abstract

Elucidating the systematic position of two Chinese species described originally as *Lactuca hirsuta* and *L. scandens*, of which only historical specimens from the late 19th and early 20th century were known, field work confirmed the occurrence of three different species. Molecular phylogenetic analysis of these species based on sequences of the nuclear ribosomal internal transcribed spacer (*nrITS*) region uncovered a hitherto unknown lineage in a phylogenetic backbone of the subtribe Crepidinae of the sunflower family tribe Cichorieae. Substantiated by comparative morphological studies, this lineage is described as genus new to science, named *Sinoseris*, endemic to the Chinese provinces Sichuan and Yunnan. Two of its three species are new to science, while the third is conspecific with both *L. hirsuta* and *L. scandens*.

## Introduction

Investigations into the flora of China have considerably increased our knowledge of the various plant groups in this country during the last decades and they have also pointed out knowledge gaps including still insufficiently known taxa. This also applies to the members of tribe Cichorieae of the largest flowering plant family Asteraceae (sunflower family). One of the least known species in this tribe was described as *Lactuca hirsuta* Franch. (Franchet, 1895) from Yunnan and has been known so far from only a few collections of the late 19th and early 20th century. Shih (1997) assigned it to *Chaetoseris*, which was later found to be a congener of *Melanoseris* (Shih & Kilian, 2011; Wang et al., 2013; Kilian et al., 2017). Doubts on this systematic position of *L. hirsuta* were already expressed by Shih & Kilian (2011). In the context of the general poverty of morphological features coupled with extensive parallel evolution in this tribe, which renders the recognition of lineages difficult (Kilian et al., 2009a), the scarcity of fruiting material available for this taxon and the absence of any recent collection of it so far have hindered re-addressing its systematic position.

The situation changed when the first author discovered new specimens that matched the description and historical specimens of *L. hirsuta* from Dayao County collected in the frame of the fourth national survey of traditional Chinese medicine resources. More in-dept studies, including special field trips guided by the re-evaluated historical collections disclosed the involvement of another taxon, described as *Lactuca scandens* C. C. Chang (1934) and not so far identified with any of the known species (Shih & Kilian, 2011), and brought to light that actually a separate lineage of three species, two of them hitherto unknown to science, is involved.

The aims of the study presented in this paper were to test the previous hypotheses of the systematic position of this taxon and its allies by comparatively studying their morphology and including them in a molecular phylogenetic analysis, and finally to draw the taxonomic conclusions from the phylogenetic evidence.

## Material and methods

### Plant material

The study is based on life plants observed and documented in the field during two field trips made to Sichuan and Yunnan in 2017 and 2018 as well as on newly collected and historical specimens of *Lactuca hirsuta* and its allies. Historical specimens were studied from the herbaria of GH, LBG, P, PE and W. Newly collected specimens were deposited in KUN, with duplicates in B and PE. For comparative morphological studies of *L. hirsuta* and allies on one hand and related genera on the other hand the herbarium collections in B, KUN and PE were consulted.

### Morphological studies

For scanning electron microscopy, achenes and pollens were directly mounted onto SEM stubs on double-sided sticky tape, coated with 20 nm Pt-Pd using a Cressington 108Auto sputter-coater and examined using a ZEISS SIGMA 300.

### Sampling, DNA extraction, amplification, sequencing and phylogenetic analysis

Multiple samples of all three species were sequenced and included in the molecular phylogenetic analysis (see Table 1) based on the nuclear ribosomal internal transcribed spacer (*nrITS*) using otherwise already published sequences. The sampling was designed to represent all genera of the Crepidinae (except for the extremely rare monotypic *Spiroseris* of Pakistan of which no sequence data are available) and its sister subtribe Chondrillinae (see Kilian et al., 2009b+; for the up-do-date systematics). *Launaea sarmentosa* (Hyoseridinae) as used as outgroup and root of the phylogenetic trees. INSDC (International Nucleotide Sequence Database Collaboration) accession numbers of published sequences follow the taxon name in the tree (Fig. 1). Extraction of DNA and amplification and sequencing of the *nrITS* region was done as described by Wang et al. (2013). Sequences were aligned with MAFFT version 7 using default parameters (Katoh et al., 2013) and the alignment adjusted manually using PhyDE version 0.9971 (Müller et al., 2010). Indels were coded as binary characters using simple indel coding (Simmons & Ochoterena, 2000) implemented in SeqState v.1.40 (Müller, 2005a). The matrix was subdivided into four partitions: ITS1, 5.8s, ITS2, indels. Phylogenetic relationships were reconstructed using maximum parsimony (MP), maximum likelihood (ML) and Bayesian inference (BI). MP was performed with the parsimony ratchet using PRAP v.2.0 (Müller, 2004) with default parameters in combination with PAUP v.4.0b10 (Swofford, 2003); Jackknife (JK) support values were calculated in PAUP with 10,000 jackknife replicates using the TBR branch swapping algorithm with 36.788% of characters deleted and one tree held during each replicate. ML analyses were done with RaxML (Stamatakis, 2014) in the version of RAxMLHPC v.8 on XSEDE on the CIPRES Science Gateway (Miller et al., 2010), using rapid bootstrapping integrated with the ML search for the optimal tree applying the general time-reversible (GTR) + Γ model. The BI analyses were performed with the MPI version of MrBayes (Ronquist et al., 2012) on the Soroban high-performance computing system of the Freie Universität Berlin. Instead of apriori testing, the optimal substitution model was sampled across the entire general time reversible (GTR) model space in the Bayesian MCMC analysis (Huelsenbeck et al., 2004). Two simultaneous runs of four parallel chains each were performed for 3 × 10^7^ generations with a sample frequency of 1 tree per 2000 generations. Convergence of the runs was checked by making sure that the average standard deviation of split frequencies of the post-burn-in runs was below 0.01 and the effective sampling size (ESS) well above 200 in either run for all parameters. TreeGraph v.2 (Stöver & Müller, 2010) was used to visualise the trees with statistical node support.

**Table 1.** The specimens’ information and accession number of newly generated sequences

| Sequence name | Collector & Collection no. (herbarium) | Locality | Accesssion no. |
| --- | --- | --- | --- |
| *Acanthocephalus benthamianus* DB464 | L. Martins & T. Janssen 1005 (JE) | Kirgistan, Jalal-Abad Oblast, Bazar-Korgon Rayon, Dscharadar |  |
| *Chondrilla canescens* DB310 | K. H. Rechinger 62177 (B100027072) | India, Kashmir, Karawring S Srinagar versus Chari Sharif |  |
| *Chondrilla leiosperma* DB306 | T. Dürbye 1754 (B100096856) | Kyrgyzstan, Tien-Shan, Issyk-Kul-See, Ak-Sakji, |  |
| *Chondrilla juncea* DB303/2005 | R. Willing & E. Willing 119544 (B100142299) | Greece, Nomos Korinthias, N Galatas |  |
| *Chondrilla chondrilloides* DB309 | Wraber 102782 (B100138369) | Italy, Udine, prope Venzona |  |
| *Heteroderis pusilla* NK131 | K. H. Rechinger 35031 (B) | Afghanistan, Tirin |  |
| *Sinoseris changii* 1 | Wang Zehuan & Chen Yaping WZH20171001 (KUN) | Yunnan Province, Dayao County, Tanhua Village |  |
| *Sinoseris changii* 2 | Wang Zehuan & Chen Yaping WZH20171011 (KUN) | Yunnan Province, Dayao County, Guihua Village |  |
| *Sinoseris changii* 3 | Wang Zehuan & Chen Yaping WZH20171018 (KUN) | Yunnan Province, Yongren County, Yongxing Village |  |
| *Sinoseris scandens* 1 | Wang Zehuan & Chen Yaping WZH20171019 (KUN) | Sichuan Province, Yanbian County, Gesala Village |  |
| *Sinoseris scandens* 2 |  |  |  |
| *Sinoseris scandens* 3 |  |  |  |
| *Sinoseris triflora* 1 | Wang Zehuan & Li Huimin WZH20181001 (KUN) | Sichuan Province, Muli County, Xiamaidi Village |  |
| *Sinoseris triflora* 2 |  |  |  |
| *Sinoseris triflora* 3 |  |  |  |
| *Syncalathium disciforme* NK168 | T. N. Ho et al. 1196 (CAS 919951) | China, Qinghai, 4300 m |  |

## Results

The *nrITS* phylogeny (Fig. 1) places the species originally described as *Lactuca hirsuta* and its two allies separate from all other genera into a well-supported clade (JS = 98.1, PP = 1, BS = 70) of the large, chiefly E Asian *Dubyaea-Nabalus-Soroseris-Syncalathium* polytomy. Finding the three species nested in one clade agrees with the conspicuous overall morphological similarity of them. The sister group relationship (with full statistical support) of two of them (Fig. 1 as *Sinoseris scandens* and *S. triflora*), also corresponds with the stronger morphological similarity between these two species compared to the third (Fig. 1, as *S. changii*), which is apparent in particular with respect to achene morphology and the larger capitula.

The three species of this clade are also morphologically clearly distinct from the other members of the *Dubyaea-Nabalus-Soroseris-Syncalathium* polytomy. Diagnostic for this clade is a synflorescence of secund, subspiciform to paniculiform paracladia, the involucre with very few, inconspicuous outer phyllaries, beaked achenes with more than two secondary ribs per main rib and a dirty white to pale brownish pappus of moderately coarse, scabrid bristles (Table 2).

**Table 2.** Comparison of diagnostic morphological features between the genera of the *Dubyaea-Nabalus-Soroseris-Syncalathium* clade and *Sinoseris* (compare the Crepidinae backbone, Fig. 1).

| Genus | Life form | Division of leaves<sup>†</sup> | Indumentum of leaves and stems | Synflorescence | Differentiation of involucre<sup>‡</sup> | Outer phyllaries of involucre | Corolla colour<sup>§</sup> | Achene apex<sup>¶</sup> | Achene ribbing pattern: main ribs /secondary ribs per main rib | Achene indumentum | Pappus colour |
| --- | --- | --- | --- | --- | --- | --- | --- | --- | --- | --- | --- |
| *Dubyaea* | perennial | UD to PD | mostly hispid, glandular or eglandular, more rarely none | corymbiform, rarely umbelliform, or capitula solitary | I to D | several, imbricate | Y or C | T | 5 /≥2 | none (or with short appressed papillae) | yellowish, brownish, rarely white |
| *Hololeion* | rhizomatous perennial | UD | none | paniculiform to corymbiform | D | several, imbricate | Y | T | 5 weak/ indistinct | none | straw-coloured |
| *Nabulus* s.l. | perennial | UD to PD | ±tomentose, or more rarely hispidulose, scabrid or villose, very rarely none | paniculiform, racemiform or spiciform | D | several, imbricate | Y or W or C | T | 5/≥ 2 | none | straw-coloured to brown |
| *Sonchella* | perennial | UD | ±none | racemiform or paniculiform | D | several, imbricate | Y | T | 5/1-2 | with short appressed papillae | white |
| *Sorosieris* | perennial | UD to PD | none to pilose | syncalathia | D | 2 | Y or W | T, rarely B | 5/ 1 or ≥ 2 | none or with short appressed to spreading erect papillae | whitish to straw-coloured, distally often greyish |
| *Syncalathium* | perennial | UD to PD | none or villous | syncalathia | U | 0 | Y or C | T | 5 /≥2 | none | greyish white |
| *“Youngia racemifera”* | perennial | UD | none | racemiform, secund | D | several, imbricate | Y | T | 5/2-3 | none | straw coloured to pale brown |
| *Sinosieris* | Annual (to monocarpic biennial) | UD to LD | conspicuously hirsute, eglandular | paniculiform of subspiciform to paniculiform paracladia | D | few, very small, inconspicuous | Y | B | 5/≥ 2 | with appressed to (apically) spreading-erect linear flattened acute papillae | dirty white to pale brownish |

†: UD, Undivided. PD, Pinnately divided. LD, Lyrately pinnatisect. ‡: I, Imbricate. D, Differentiated in inner and outer series. U, Uniseriate. §: Y, Yellow. C, Cyanic. W, White. ¶: B, Beaked, T, Truncate.

Consequently, this lineage is best classified as an own genus, for which we have chosen the name *Sinoseris* (see Taxonomy, below). The genus is endemic to China, where its species are restricted to Yunnan and Sichuan (Fig. 2). Only the most widespread of the three species has been known to science so far. Its original name *Lactuca hirsuta* is illegitimate as a younger homonym of a name coined much earlier for a species of North America. The Chinese species was described a second time 40 years later by Chao Chien Chang (1900–1972) as *L. scandens*, a name that has to be taken now as basionym for this species. The conspecifity of both taxa was blurred, however, by infraspecific variation and incorrectly described features in the protologue of the first name (see Taxonomy, below). The second species (see Taxonomy below, under *S. triflora*), hitherto undescribed, was discovered by us first among the historical specimens from the early 20th century determined as *L. hirsuta* before we succeeded to recollect it from the wild. The third species (*S. changii*) was not known to us from historical herbarium material, but ITS sequences published under the name *L. scandens* (INSDC acc. no. KF732051 to KF732056, see Fig. 1) and included in our phylogenetic reconstruction were found nested in the same clade as our sequences of this undescribed species.

## Discussion

The Asian-North American *Dubyaea-Nabalus-Soroseris-Syncalathium* polytomy, in which the clade with the species under study is nested, is one of six major terminal clades resolved in our phylogenetic backbone of the subtribe Crepidinae spanning all genera (Fig. 1). This seems surprising in view of their previous classification as members of *Lactuca* or the Lactucinae. It must be considered, however, that the generic concept of *Lactuca* in the late 19th and early 20th century was extremely wide and spanned members of several of the modern subtribes (Kilian et al., 2017). Disentangling the subtribes Lactucinae and Crepidinae has, moreover, proven particularly difficult due to the scarcity of non-homoplastic morphological synapomorphies (Bremer, 1994).

The *Dubyaea-Nabalus-Soroseris-Syncalathium* clade has been resolved in several studies with varying partial representation of its members (Kilian et al., 2009a; Zhang et al., 2011; Liu et al., 2013; Kilian et al., 2017). Estimations divergence times for the Crepidinae by Zhang et al. (2011) revealed that the *Dubyaea-Nabalus-Soroseris-Syncalathium* clade is likely of Pliocene origin, with a crown age of around 5 myr only; for genera such as *Soroseris* or *Syncalathium* a crown age of less than 2 and 3 myr, respectively, have been estimated. The comparatively young age of this clade may be responsible for the shallow genetic differentiation among many of its members (Zhang et al., 2011), in particular within lineages such as *Soroseris* or the *Nabalus trifoliatus* lineage, as indicated by branch lengths in Fig. 1. The comparatively deep genetic differentiation between the first sister clades of *Sinoseris* is all the more conspicuous.

All three known species of the clade classified here as the new genus *Sinoseris* are annuals (to, perhaps, monocarpic biennials) and grow in the subtropical (warm-temperate) highlands of Sichuan and Yunnan, where they all seem to grow preferably in open rocky habitats. All other members of the *Dubyaea-Nabalus-Soroseris-Syncalathium* clade are, in contrast, perennials and confined to the higher montane or alpine zone (*Dubyaea, Soroseris, Syncalathium, “Youngia “ racemifera*) or more continental and either colder (*Nabalus, Hololeion*) or more arid (*Sonchella*) climates. The *Sinoseris* lineage is therefore somewhat outstanding in this clade, which deserves further attention.

The species have been rarely collected so far, which appears surprising as they are not inconspicuous. Reasons may be their late flowering from late September onwards and their scattered occurrence, perhaps due to their preference of rocky habitats.

### Taxonomy

***Sinoseris*** N. Kilian, Ze H. Wang & H. Peng, **gen. nov**.

Type: *Sinoseris scandens* (C. C. Chang) Ze H. Wang, N. Kilian & H. Peng

#### Diagnostic features

Annual (to monocarpic biennial) herbs; stems, leaves and involucres with conspicuous hirsute indumentum; basal leaves distinctly petiolate; synflorescence of secund, subspiciform to paniculiform paracladia; capitula with 3–12 florets; involucre with inconspicuous outer phyllaries; achenes beaked; achene corpus with 5 main ribs alternating with (2–)3–4 secondary ribs; pappus dirty white to pale brown of moderately coarse scabrid bristles.

#### Description

*Annual (to monocarpic biennial?) herbs* with leafy stem and conspicuous hirsute indumentum on stem, leaves and involucres, late flowering (Sep-Nov). *Basal and lowermost cauline leaves* with petiole-like portion as long as or longer than lamina. *Capitula* 3–12 flowered, in ± secund, subspiciform to narrowly paniculiform paracladia from the axils of the cauline leaves, pendent in bud, subpendent at anthesis and pendent again at fruiting. *Involucre* narrowly cylindrical at anthesis, strongly differentiated into an equal inner and very inconspicuous outer phyllary series. *Receptacle* epaleate, glabrous and smooth. *Florets* with yellow corolla, styles greyish to blackish. *Pollen* echinolophate, tricolpate, of the *Cichorium* type (sensu Blackmore 1986) with polar areas either triangular, moderately extensive and each with c. 12 spines, or very extensive, approximately hexagonal and each with >20 spines, and with moderately narrow interlacunar gaps (Fig. 4e–f, 6e–f, 8e–f). *Achenes* beaked; corpus somewhat flattened, ± subconical, with 5 main ribs (best discernable near base) alternating with (2–)3–4 secondary ribs (fully developed in middle third and then similar in shape to main ribs) dark brown, with appressed to spreading-erect (in distal portion of corpus) linear flattened acute papillae; beak slender, shorter than or as long as corpus. *Pappus* dirty white, caducous, of scabrid bristles similar in length and diameter, with 7–12(–14) rows of cells in cross section near base.

#### Etymology

The generic name is a compound of the Latin name “Sina” for China and “seris” (σέρις;), the classical Greek name for salad (more precisely of *Cichorium* species).

### Key to the species of *Sinoseris*

1 Involucre with usually 8 inner phyllaries; capitula with 8–12 florets; anther tube golden yellow to brownish; achenes abruptly contracted into a slender beak as long as the obconical and below the beak broad-shouldered corpus………………***S. changii***

1 Involucre with 5 or less inner phyllaries and 6 or less florets; anther tube blackish; achenes attenuate into a slender beak much shorter than the narrowly ellipsoidal corpus…………**2**

2 Involucre with (4–)5 inner phyllaries; capitula with 4–6 florets ………***S. scandens***

2 Involucre with 3 inner phyllaries; capitula with 3 florets………***S. triflora***

### 1. *Sinoseris scandens* (C. C. Chang) Ze H. Wang, N. Kilian & H. Peng, comb. nov

≡ *Lactuca scandens* C. C. Chang in Contr. Biol. Lab. Sci. Soc. China, Bot. Ser. 9: 133. 1934 – Holotype: [China, Sichuan] “Yien-Pien Hsian” [not “vicinity of Pa-hsien (Chungking)”] [c. 26.90°N, 101.56°E], Oct 1932, *T T. Yü 1702* (**lectotype selected here:** LBG00092880! [marked as “holotype”]; isolectotypes: LBG00092879!, PE01106722! [marked as “isotype”]).

= *Lactuca hirsuta* Franch. in J. Bot. (Morot) 9: 258. 1895 [non *Lactuca hirsuta* Nutt. 1818] ≡ *Chaetoseris hirsuta* C. Shih, Fl. Reipubl. Popularis Sin. 80(1): 282. 1997 ≡ *Melanoseris hirsuta* (C. Shih) N. Kilian in Wu et al., Fl. China 20–21: 220. 2011. – Syntypes: China, Yunnan, lieux ombragés au mont Che-tscho-tze, au-dessus de Tapintze [a village at c. 26.10°N, 100.04°E], 10 Oct 1882, *J. M. Delavay 627* (**lectotype selected here:** P00288022! [marked as “holotype”]; isolectotype P00750292! [marked as “isotype”]).

#### Description

*Annual (to monocarpic biennial) herbs*, 15–80 cm tall, strongly hirsute of eglandular reddish-purplish hairs, with a taproot. *Stem* solitary, or if branched right from the base, plants seemingly with several stems, erect or arched erect, branching, leafy. *Basal and lower cauline leaves* abruptly contracted into a petiole-like portion up to 15 cm long, its base semi-amplexicaul to, more often, winged and distinctly clasping the stem; lamina broadly triangular, ovate to broadly lanceolate, or oblanceolate, 3–14 cm long, 2.5–13.5 cm wide, entire to lyrately pinnatisect, with a broadly ovate to broadly triangular terminal lobe, cordate or obtuse to cuneate at base and acute at apex, and with 1–2 pairs of much smaller triangular to rhombic, acute to obtuse, or up to 5 pairs of very small ovate to elliptic lateral lobes; lamina margin variably shallowly or deeply sinuate-dentate, often irregularly so, and denticulate. *Middle and upper cauline leaves* oblanceolate or ovate to lanceolate, smaller, with winged or ± without petiole-like basal portion, otherwise similar to lower cauline leaves, base distinctly clasping. *Synflorescence* of a flowering stem in well-developed plants with several paracladia from the axils of the cauline leaves, all subspiciform to narrowly paniculiform and ± secund, each with a few to more than a dozen capitula pendent in bud, subpendent at anthesis and pendent again at fruiting. *Capitula* with (4–)5(–6) florets; peduncle in most cases less than 1 cm long. *Involucre* narrowly cylindrical, 10–13 mm long; strongly differentiated into inner and outer phyllary series, the latter even very inconspicuous; phyllaries abaxially reddish hirsute as remainder of the plant; outer phyllaries up to 3, very inconspicuous, narrowly linear, 0.9 mm × 0.2 mm; inner phyllaries usually 5, linear-lanceolate and similar in length, green, sometimes (partly) with a purplish tinge. *Receptacle* epaleate, glabrous and smooth. *Florets* with bright yellow corolla; ligule broadly elliptical to obovate, ± horizontally spread, 14–16 mm long and up to 3 mm wide, tube c. 6 mm long; anther tube blackish, fertile part 4.6–4.8 mm long, apical appendages rounded, c. 0.2 mm long, basal appendages c. 0.6 mm long; *style* and style arms blackish. Pollen of the *Cichorium* type (sensu Blackmore 1986) with triangular, moderately extensive polar areas each with c. 12 spines and with moderately narrow interlacunar gaps (Fig. 4e–f). *Achenes* 8–9 mm long, corpus narrowly ellipsoidal to subconical, subcompressed, dark brown mottled white, covered with linear flattened acute antrorse papillae, shorter and appressed in the lower two thirds of the corpus, longer and spreading-erect in the upper third, with 5 main ribs (best discernible near base) alternating with (2–)3(–4) secondary ribs (fully developed in middle third and then similar in shape to main ribs), apex attenuate into a slender whitish beak of 3–4 mm (Fig. 4a–d). *Pappus* c. 6–7 mm long, dirty white to pale brownish, caducous, bristles of similar length and diameter, near base of 8–14 rows of cells in cross section.

#### Distribution

NW Yunnan and SW Sichuan.

#### Habitat and ecology

The species occurs at altitudes between 1750 m and c. 3000 m on rocky slopes with open grassland vegetation. Fl. and fr. Sep–Nov.

#### Notes on typification and synonymy

The description in the protologue is detailed and well corresponds to the type material. The eglandular hirsute reddish indumentum in combination with capitula of (mostly) 4 or 5 yellow florets, blackish anther tubes and styles, and an involucre of (4 or) 5 inner and very inconspicuous outer phyllaries diagnose the taxon perfectly. The author compares the species with *L. hirsuta* Franch. as its “nearest ally”, stating that *L. scandens* differs from that by a red (versus “white”, actually “dirty white” [“setis sordidis”]) hispid eglandular (versus mixed eglandular-glandular) indumentum, a “white” (versus “dirty white”) pappus, and undivided (versus lyrately pinnate) leaves. The description of the pappus of *L. scandens* is contradicting in the protologue: the Latin description states “setae … sordide albae”, thus “dirty white” instead of “white”, which agrees with our observation and also with the description of *L. hirsuta*. The leaves in *L. hirsuta* are lyrately pinnatisect but they are variable within populations, ranging from entire to pinnatisect. The indumentum in the type specimens of *L. hirsuta* looks in fact dirty white but this is likely only an effect of drying; the statement about presence of glandular hairs in the protologue of *L. hirsuta* seems erroneous. Also the number of 8 florets per capitulum given in the protologue of *L. hirsuta* is apparently erroneous. With the help of the curators in P their number has been confirmed as 5–6. The number of inner phyllaries is correctly and in agreement with *L. scandens* given as 5. It can safely be concluded that both taxa are conspecific.

The gathering of Pére Jean Marie Delavay was made in the mountains above the village “Tapintze” [= Dapingzi], where Père Delavay lived from August 1882 onwards for some years as missionary. The village is situated at c. 26.10°N, 100.04°E, some 65 km NE of Dali in the valley Loulou river, a tributary of the Yangtze, which passes c. 15 km NE (Kilpatrick, 2014: 69, 102; reproducing a map of the area drawn by P. Delavay). Technically the two sheets of the same gathering by Père Delavay of *Lactuca hirsuta* Franch. at P are syntypes so that a lectotype has to be designated. We have designated the one marked as “holotype”, with the original label of the collector and the determination in Franchet’s hand.

#### Further specimens seen

China: Setchuan, in regio subtropica convallis fluminis Yalung ad affluentem versus Yenyüen in altograminetis ad viam Gwanyingai, 27.33°N [c. 102°E], subtr. calceo, 1750 m, 29 Sep 1914, *H. Handel-Mazzetti 5342* [Diar. Nr. 887nota] (W 1940–348!)

Setchuan, in montis Lungdschu-schan prope urbem Huili regione temperata, rupibus supra vicum Djindjiatsun in limite reg. calide temperatae, 2800 m, [c. 26.75°N, 102.23°E], 16 Sep 1914, *H. Handel-Mazzetti 5186* [Diar. Nr. 839] (W 1940–347!)

Sichuan, Panzhihua, Yanbian County, Gesala Village, 27.130136°N, 101.287422°E, elev. 2523 m, 14 October 2017, *Wang Zehuan & Chen Yaping WZH20171019* (B!, KUN!, PE!)

Yunnan, Lijiang, Ninglang Yi Autonomous County, from Zhanhe Village to Yongningping Village, 26.742985°N, 101.000726°E, elev. 2934 m, 20 Oct 2018, *Wang Zehuan & Li Huimin WZH20181005* (KUN!)

#### Threat status

*Sinoseris scandens* has been collected so far from five separate localities in Sichuan and Yunnan. Three of them are only known from historical collections made between 1882 and 1932. At the two current localities, at Gesala <500 individuals have been counted in 2017, and at Ninglang <100 individuals have been counted in 2018. Population sizes seem to vary, however, considerably between the years as observed in Gesala comparing 2017 and 2018. A formal threat status assessment according to the IUCN criteria and categories (IUCN 2001) for this species requires more data on its actual and historical distribution and population sizes in relation to land use changes. Although the scarcity of collections of this species may be partly due to its late flowering in the year, *S. scandens* seems to have a scattered distribution and be a rare species endemic to some small part of the two provinces. Its status should be thus of concern and addressed by further investigations.

### 2. *Sinoseris triflora* Ze H. Wang, N. Kilian & H. Peng, sp. nov

#### Holotype

China, Sichuan, Yi Autonomous Prefecture of Liangshan, Muli Tibetan Autonomous County, Xiamaidi Village, from Mianbu bealock to Muli County, 27.730014°N, 101.236871°E, elev. 3022 m, 19 Oct 2018, *Wang Zehuan & Li Huimin WZH20181001* (KUN!, isotypes: B!, KUN!, PE!)

#### Diagnosis

The species can be easily distinguished from the other two species by involucre with 3 inner phyllaries, capitula with 3(–4) florets and 3(–4) achenes.

#### Description

*Annual (to monocarpic biennial) herbs* with taproot, 15–90 cm tall, strongly hirsute of eglandular pale to dark reddish-purplish hairs. *Stem* solitary, erect, branching from base or higher up, leafy. *Basal and lower cauline leaves* abruptly contracted into a petiole-like portion up to 6 cm long, its base semiamplexicaul or winged and distinctly clasping the stem; lamina orbicular (in particular in basal leaves) to broadly triangular or ovate, or oblanceolate, 4–8 cm long, 2.8–5.5 cm wide, entire or lyrately pinnatifid to pinnatisect, with a large orbicular to broadly triangular terminal lobe with cordate or obtuse to cuneate base and acute apex, and one to several pairs of (much) smaller ovate or elliptic to rhombic or ± triangular, acute to obtuse lateral lobes; lamina margin variably shallowly or deeply sinuate-dentate, often irregularly so, and denticulate. *Middle and upper cauline leaves* ovate to lanceolate, smaller, with winged or ± without petiole-like basal portion, otherwise similar to lower cauline leaves, base distinctly clasping the stem. *Synflorescence* of a flowering stem in well-developed plants with several paracladia from the axils of the cauline leaves, all subspiciform to narrowly paniculiform and ± secund, each with a few to more than a dozen capitula pendent in bud, subpendent at anthesis and pendent again at fruiting. *Capitula* with 3 florets; peduncles mostly shorter than the involucre. *Involucre* narrowly cylindrical, 10–12 mm long; strongly differentiated into inner and ± inconspicuous outer phyllary series; phyllaries abaxially pale to reddish hirsute as remainder of the plant; outer phyllaries 2, narrowly linear, 2–4 × 0.2–0.3 mm; inner phyllaries 3, linear-lanceolate and similar in length, green, sometimes (partly) with a purplish tinge. *Receptacle* epaleate, glabrous and smooth. *Florets* with yellow corolla; ligule elliptical, reflexed, 9–12 mm long and up to c. 2 mm wide, dorsally pale yellow, tube c. 5–6 mm long; anther tube blackish, fertile part 2.8–3.6 mm long, apical appendages rounded, c. 0.2 mm long, basal appendages c. 0.3 mm long; *style* and style arms blackish. Pollen of the *Cichorium* type (sensu Blackmore 1986) with very extensive, approximately hexagonal polar areas, each with >20 spines, and with moderately narrow interlacunar gaps (Fig. 6e–f). *Achenes* 7–9 mm long, corpus subconical, with 5 main ribs (best discernable near base) alternating with 3(–4) secondary ribs (fully developed in middle third and then similar in shape to main ribs), dark brown, with linear flattened acute antrose papillae, shorter and appressed in the lower two thirds of the corpus, longer and spreading-erect in the upper third; apex of corpus attenuate into a slender whitish beak of c. 1–2 mm (Fig. 6a–d). *Pappus* c. 6–7 mm long, dirty white, caducous, bristles of similar length and diameter, near base of 8–12 rows of cells in cross section.

#### Distribution

E Yunnan and SW Sichuan.

#### Habitat and ecology

The species grown on rocky stream banks and slopes with open bushy vegetation at altitudes between c. 2200 and 3250 m. Fl. and fr. Sep–Nov.

#### Further specimens seen

China: Setchuan, inter oppidum Yenyüen et castellum Kwapi [“between the cities Yanyuan and the fortress ‘Kwapi’”, name not traced], c. 27.75°N [c.101.55°E, very rough estimate], ad viam vico Tangetu oppositam [“at the road opposite the village Tangetu”, place not traced], substr. arenaceo, 3250 m, 4 Oct 1914, *H. Handel-Mazzetti 5476* [Diar. Nr. 923] (GH!, W 1940–349!)

Yunnan, Lou-Pou, prefecture de Tong Tchouan [= Dongchuan, city = 26.082872°N, 103.18783°E], 19 Oct 1906, *F. Ducloux 4477* (P3732867!, P03732866! [label text: “Yunnan, Lou Pou à deux journees de Tong Tchouan, plante cueillie par Joseph Tschang, 19 Oct 1906, *F. Ducloux 4477”]*)

Sichuan, Yi Autonomous Prefecture of Liangshan, Muli Tibetan Autonomous County, Xiamaidi Village, from Mianbu bealock to Muli County, 27.730082°N, 101.236879°E, elev. 3021 m, 11 Oct 2018, *Chen Yaping, Jiang Lei & Zheng Hailei EM652* (KUN!).

Sichuan, Yi Autonomous Prefecture of Liangshan, Muli Tibetan Autonomous County, Sanjiaoya Town, on the way from Biji to Guoquanyan, 28.107632°N, 101.467566°E, elev. 2275 m, 13 Oct 2018, *Chen Yaping, Jiang Lei & Zheng Hailei EM672* (KUN!).

#### Threat status

The species is known from two historical and three current localities. At these three populations in the Muli County only 10, 80 and 250 individuals were counted respectively, so that the population sizes seem in general smaller than the other two species. The species occurs in an only scattered way in its distribution area, similar to *S. scandens*, which may be due to its requirement of open rocky habitats. Nearly a third of the few-flowered capitula of many individuals was found infected in 2018 by some insects species and did therefore not produce fruits. The species is certainly rare and of localised distribution. Its status should be thus of concern and addressed by further investigations.

### 3. *Sinoseris changii* Ze H. Wang, N. Kilian & H. Peng, sp. nov

#### Holotype

CHINA. Yunnan: Chuxiong Autonomous Prefecture, Dayao County, Tanhua Village, the mountain behind Tanhua Temple, 25.950745°N, 101.232407°E, elev. 2719 m, 12 Oct 2017, Wang Zehuan & Chen Yaping *WZH20171001* (KUN!, isotypes: B!, KUN!, PE!).

#### Diagnosis

The species can be easily distinguished from the other two species by involucre with usually 8 inner phyllaries; capitula with 8–12 florets and 8–12 achenes; anther tube golden yellow to brownish; achenes abruptly contracted into a slender beak as long as the obconical and below the beak broad-shouldered corpus.

#### Description

*Annual (to monocarpic biennial) herbs* with taproot, 20–100 cm tall, strongly hirsute of eglandular pale to dark reddish-purplish hairs. *Stem* solitary, or if branched right from the base, plants seemingly with several stems, erect, branching, leafy. *Basal, lower and middle cauline leaves* abruptly contracted into a petiole-like portion up to 17 cm long with 0–3 pair(s) of lobes, otherwise unwinged, and with semi-amplexicaul to at most weakly clasping base; lamina rhombic, ovate, triangular or lanceolate in outline, 3–11 cm long, 3–8 cm wide, entire to lyrately pinnate with 1–2(–3) pair(s) of smaller acute or obtuse lateral lobes and a large terminal lobe, fresh green on upper and paler, greyish green on the lower face; margin irregularly sinuate-dentate and denticulate; apex ± acute, base distinctly cordate. *Upper cauline leaves* similar to middle ones but smaller, or ± oblanceolate and the entire or lyrately pinnatifid lamina attenuate into a petiole-like portion much shorter than the lamina, or ± lanceolate and sessile with semi-amplexicaul to weakly clasping base. *Synflorescence* of a flowering stem paniculiform to corymbiform of some to many capitula erect in bud, spreading-erect to subpendent at anthesis and pendent at fruiting; in well-developed plants with several paracladia from the axils of the cauline leaves. *Capitula* with 8–12 florets; peduncles mostly longer than the involucre. *Involucre* narrowly cylindrical, 9–14 mm long; strongly differentiated into inner and ± inconspicuous outer phyllary series; phyllaries abaxially pale to reddish hirsute as remainder of the plant; outer phyllaries narrowly linear, 2–5, 0.7–1.9 × 0.3 mm; inner phyllaries usually 8, linear-lanceolate and similar in length, green, sometimes (partly) with a purplish tinge. *Receptacle* epaleate, glabrous and smooth. *Florets* with yellow corolla; ligule broadly elliptical to obovate, ± horizontally spread, 10–12 mm long and up to 3 mm wide, dorsally pale yellow, tube 5–6 mm long; anther tube golden yellow to brownish, fertile part 2.6–2.8 mm long, apical appendages rounded, c. 0.3 mm long, basal appendages c. 0.5 mm long; *style* and style arms pale greyish to blackish. *Pollen* of the *Cichorium* type (sensu Blackmore 1986) with very extensive, approximately hexagonal polar areas, each with >20 spines, and with moderately narrow interlacunar gaps (Fig. 8e–f). *Achenes* 6–7 mm long, corpus subconical, with 5 main ribs (best discernible near base) alternating with 3(–4) secondary ribs (fully developed in middle third and then similar in shape to main ribs), dark brown, with linear flattened antrose papillae, spreading-erect at the apex and appressed and shorter in the rest, at its widest diameter abruptly contracted into a slender pale brown, basally appressed papillate beak of about the same length as the corpus, corpus below the beak broad-shouldered (Fig. 8a–d). *Pappus* 5–6 mm long, dirty white to pale brownish, caducous, bristles of similar length and diameter, near base of 7–12 rows of cells in cross section.

#### Distribution

Central Yunnan

#### Habitat and ecology

The species is confined to open rocky habitats. Fl. and fr. Sep–Oct.

#### Etymology

We dedicate this species to the memory of Chao Chien Chang (1900–1972), one of the early modern Chinese botanists. He worked at the Kunming Institute of Botany of the Chinese Academy of Sciences, studied on Chinese Asteraceae, in particular these taxa distributed in Yunnan.

#### Further specimens seen

CHINA. Yunnan: Chuxiong Autonomous Prefecture, Dayao County, Wanbi Village, Gao-ping-zi, 26.224444°N, 101.308611°E, elev. 2229 m, 15 Oct 2015, Exped. Dayao team *ly334* (KUN!). CHINA. Yunnan: Chuxiong Autonomous Prefecture, Dayao County, Tanhua Temple to Guihua Village, 26.042925°N, 101.273300°E, elev. 2103 m, 12 Oct 2017, *Wang Zehuan & Chen Yaping WZH20171003* (KUN!).

CHINA. Yunnan: Chuxiong Autonomous Prefecture, Dayao County, Tanhua Temple to Guihua Village, 26.074892°N, 101.284225°E, elev. 1945 m, 12 Oct 2017, Wang Zehuan & Chen Yaping *WZH20171004* (KUN!).

CHINA. Yunnan: Chuxiong Autonomous Prefecture, Dayao County, Guihua Village, 26.082596°N, 101.284981°E, elev. 2719 m, 13 Oct 2017, Wang Zehuan & Chen Yaping *WZH20171011* (KUN!). CHINA. Yunnan: Chuxiong Autonomous Prefecture, Yongren County, Yongxing Village, 26.344374°N, 101.614891°E, elev. 2123 m, 13 Oct 2017, Wang Zehuan & Chen Yaping *WZH20171018* (KUN!).

#### Threat status

The occurrence of this species is less scattered and rare compared to the other two species. Between Tanhua and Wanbi, for example, the species is present in sunny rocky slopes with high frequency, although the individual populations usually do not exceed a few hundred mature individuals. Considering its localised distribution in Central Yunnan, its status should be of concern and addressed by further investigations.

## Acknowledgements

The authors are grateful to the staff from the herbarium of LBG and P for their warm help in providing related photos of type specimens. The first author also thanks the staff of KUN and Institute of cultivation and processing of Chinese medicinal materials for research facilities. This study was supported by the National Natural Science Foundation of China (grant no. 31500168).

## Figure captions

**Fig. 1.** Majority consensus phylogram of the Crepidinae from the Bayesian analysis (support values: first line: maximum parsimony jackknife, second line: Bayesian posterior probability / maximum likelihood bootstrap) based on the *nrITS* region.

**Fig. 2.** Distribution of *Sinoseris* based on specimens seen. Square, *S. scandens*. Triangle, *S. triflora*. Circle, *S. changii*.

**Fig. 3.** *Sinoseris scandens* – **a,** Habit. **b,** Flowering capitula. c, Fruiting capitula with mature achenes. **d,** Basal leaf. **e,** Sequence of stem leaves (upwards, from right to left). –Photographs by Wang Zehuan from near Sichuan, near Gesala Village on 14 Oct 2017 (a-b, d-e) and 12 Nov 2018 (c); population voucher: *Wang Zehuan & Chen Yaping WZH20171019* (KUN).

**Fig. 4.** *Sinoseris scandens* – **a–d,** Achene, apical (a) and basal (b) portion, pappus disk with detached pappus bristles showing bristles cross section(c), apex of corpus with parts of the papillae longer and spreading (d). **e–f,** Pollen, polar (e) and equatorial (f) view. – From *Wang Zehuan & Chen Yaping WZH20171019* (KUN).

**Fig. 5.** *Sinoseris triflora* – **a,** Habit. **b,** Flowering capitula. **c,** Stem leaf. **d,** Mature achene. – Photographs from Sichuan, near Xiamaidi Village by Chen Yaping on 11 Oct 2018 (a-c) and Wang Zehuan on 13 Nov 2018 (d); population voucher: *Chen Yaping, Jiang Lei & Zheng Hailei EM652* (KUN).

**Fig. 6.** *Sinoseris triflora* – **a–d,** Achene, apical (a) and basal (b) half, pappus disk with detached pappus bristles showing bristles cross section(c), apex of corpus with parts of the papillae longer and spreading (d). **e–f,** Pollen, polar (e) and equatorial (f) view. – From *Wang Zehuan & Li Huimin WZH20181001* (KUN).

**Fig. 7.** *Sinoseris changii* – **a,** Habit. **b-c,** Flowering capitula. **d-e**, Leaf indumentum. **f,** Achene. – Photographs by Chen Yaping (a) and Wang Zehuan (b-f) from Yunnan, near Tanhua Temple, on 12 Oct 2017; population voucher: *Wang Zehuan & Chen Yaping WZH20171001* (KUN).

**Fig. 8.** *Sinoseris changii* – **a–d,** Achene, apical (a) and basal (b) portion, pappus disk with detached pappus bristles showing bristles cross section(c), apex of corpus with parts of the papillae longer and spreading (d). **e–f,** Pollen, polar (e) and equatorial (f) view. – From *Wang Zehuan & Chen Yaping WZH20171001* (KUN).

